# Mapping the Structural Brain Network of Psychopathy: Convergent Evidence from Humans and Chimpanzees

**DOI:** 10.1101/2025.09.25.678605

**Authors:** Jules R. Dugré, Sam Vickery, Robert D. Latzman, William D. Hopkins, Felix Hoffstaedter, Stéphane A. De Brito

## Abstract

Psychopathy, a condition characterized by profound emotional and interpersonal deficits, has long been hypothesized to stem, in part, from structural brain abnormalities. Yet neuroimaging findings remain inconsistent. To address these discrepancies, we applied a novel structural network mapping approach combined with cross-species analyses. Traditional meta-analysis revealed weak spatial convergence across 20 samples. Nevertheless, we found that heterogeneous peak locations coalesced into a distributed set of regions encompassing the insular, and prefrontal cortices. This network overlapped strikingly (r = .93) with a lesion-derived network causally linked to antisocial behaviour, and variation within it predicted volumetric risk scores in both humans (n = 107, R² = .16) and chimpanzees (n = 148, R² = .21). These findings suggest that psychopathy reflects abnormalities across a collection of distributed regions rather than isolated areas, providing a unifying explanation for decades of inconsistent results and advancing our understanding of its clinical, biological, and evolutionary bases.

## 1. Introduction

Psychopathy is a personality disorder characterized by a constellation of symptoms such as shallow affect, pathological lying, low or lack of remorse and empathy, and antisocial behavior ^1,2^. The convention for clinically assessing psychopathy, the Psychopathy Checklist– Revised (PCL-R) ^2,3^, maps onto two overarching factors — Factor 1 (i.e., Interpersonal and Affective facets) and Factor 2 (i.e., Lifestyle and Antisocial facets) ^4^ — which can be further broken down into a four-facet structure ^5^. Approximately 1% of the general population is affected, compared to up to 25% in carceral settings ^1^. Over the past decades, the psychological and criminological correlates of psychopathy have been well described. For instance, this disorder is associated with a wide range of negative outcomes and psychosocial maladjustments, including high rates of criminal reoffending, which impose a substantial financial burden on society each year ^6^. In the general population, however, psychopathic traits have been shown to shape leadership and political affiliation ^7^. Despite its critical importance for uncovering the pathogenesis of psychopathy, neurobiological research lags behind.

Early neuroimaging research suggested that individuals with elevated psychopathic traits showed significant grey matter volume (GMV) reductions in the prefrontal cortex (PFC) ^8^. Since then, structural abnormalities in the PFC have remained one of the most consistently reported findings in the psychopathy literature ^9–13^. Indeed, the most recent meta-analysis of voxel-based morphometry (VBM) studies show significant GMV reductions in the superior frontal gyrus and medial orbitofrontal cortex (OFC) in individuals with psychopathy compared to controls ^13^. Converging evidence from lesion studies further shows that damage to the PFC ^14–17^ can produce a clinical profile resembling psychopathy, termed “*pseudopsychopathy*” ^18^ or “*acquired sociopathy*” ^17^. Similar to psychopathy, this condition involves impaired decision-making ^19,20^, reduced prosocial motivation ^21^, and diminished autonomic responses to social stimuli ^22^. While the current state of evidence suggests that structural changes to the PFC consistently contribute to antagonistic behaviors in humans, similar patterns have also been observed in non-human primates. Intriguingly, variations of GMV in PFC (but also in parietal regions) have also been reported to differentiate between chimpanzees and bonobos (bonobos > chimpanzees, ^23^), potentially accounting for their pronounced differences in propensity for aggressive behavior ^24^. Indeed, Latzman and colleagues ^25^ demonstrated that reductions in PFC volume (i.e., medial PFC, dorsal PFC, and ACC) in chimpanzees were linked to dominance and extraversion, core features of psychopathy. Taken together, these findings highlight the central role of paralimbic structures (especially frontal areas) in psychopathy ^26^, and provide a potential evolutionary perspective on psychopathy.

Recently, some authors have expressed their concerns towards the reliability of neurobiological findings in psychopathy ^27–29^. More precisely, Deming and colleagues ^28^ recently showed that 83.3lJ% of experiments on the structural alterations in the medial PFC yielded null effects. Putting in perspective, it is expected that most meta-analytic findings are driven by around 25% of the studies ^30^, given the substantial clinical and methodological differences between studies. Nevertheless, these findings underscore a major challenge that must be resolved before meaningful progress can be made in the field. A wide range of possible explanations for this lack of reproducibility has been proposed, including sample size ^31,32^, scan length ^33^, study design ^34^; analytical choices ^35,36^, measurement error ^37^, as well as heterogeneity in clinical samples, differences in psychopathy measures, and the operationalization of psychopathy as a unitary construct ^38,39^. While these factors are undeniably important, current research consistently violates basic assumptions of brain functioning, a practice that partially contributes to perpetuate the replication crisis in psychiatric neuroimaging ^40,41^. Indeed, current neuroimaging approaches consider brain regions as isolated units, with many efforts aiming at uncovering a single brain structure causal to psychiatric disorders (e.g., psychopathy). However, from such a localizationist perspective, it is difficult to explain how damages to distinct areas in the brain (i.e., amygdala and OFC) may contribute to a similar clinical profile ^17,18,42^.

A growing body of work instead advocates for viewing the brain as a complex system organized into functional modules of highly interconnected brain regions, which align with our long-lasting knowledge about the distributed organization of the brain ^41,43,44^. This shift in paradigm allows for new hypotheses of psychiatric conditions, such as *neurobiological equifinality* ^46^, where regional deficits in heterogeneous locations may give rise to the same phenotype. In line with this network approach, Darby and colleagues ^45^ demonstrated that heterogeneous brain lesions occurring prior to the onset of antisocial behaviors were functionally connected to a common neural network. More specifically, by using functional connectomes of 1,000 healthy adults, the authors mapped the functional connectivity profile of each of the 17 brain lesions causal to antisocial behaviors and revealed that the lesions shared a common overarching network, despite the heterogeneous locations of affected regions. Recently, Dugré and De Brito ^46^ extended this lesion network mapping ^47^ to a coordinate-based meta-analysis of 40 fMRI samples on psychopathy, and demonstrated high replicability across studies but also substantial similarity with the network identified across brain lesions by Darby and colleagues ^45^. Identifying robust and replicable structural imaging findings among VBM studies on psychopathy has yet to be achieved. To our knowledge, the application of this method to structural imaging remains largely limited and could provide a novel way to identify a common and replicable structural covariance network underpinning psychopathy. Indeed, morphological variations across brain regions are known to covary ^48^, and this covariance has been found to reflect underlying functional connectivity ^49^. However, structural covariance also reveals unique relationships that may not be fully captured by functional connectivity ^50^, suggesting a more complex biological meaning possibly involving shared aetiology and/or neural plasticity ^49,51–53^. While some studies have used functional connectivity to map GMV atrophy ^54^, retaining morphometric information may offer greater biological precision.

Therefore, we first aimed to conduct an updated meta-analysis of VBM studies on psychopathy (20 samples), given the low power of prior meta-analysis ^13^ (*k* = 7 samples) & ^11^ (k=9 samples). Secondly, we aimed to extend the network mapping approach to structural imaging data by testing whether the heterogeneous locations of GMV deficits reported across VBM studies on psychopathy are in fact structurally connected within a common structural covariance network. To this end, we used normative GMV data from a large sample of healthy individuals (n = 1000) to calculate correlations between the average volume of voxels within the identified regions and the volumes of voxels across the rest of the brain. Similarly to Dugré et De Brito ^46^, we first hypothesized that meta-analytic findings from a traditional approach would be driven by a few studies (< 25lJ%), which may partly explain the inconsistent findings observed in previous meta-analyses ^11,13^. In turn, we hypothesized that our method would allow us to identify a common structural covariance network across VBM studies on psychopathy, with replicability of some regions greater than 60lJ% across studies, especially in paralimbic regions.

Here, we showed for the first time that heterogeneous locations of GMV deficits reported across VBM studies on psychopathy indeed map onto common structural covariance network. We then tested the validity of this structural network in two ways. First, we tested whether this network would spatially correlate with a network of brain lesions causally linked to antisocial behaviours ^45^, demonstrate specificity relative to matched control lesions. Second, we conducted external validation analyses using cross-species neuroimaging data and psychopathic traits from both humans and chimpanzees.

## 2. Results

### 2.1. Included Studies

A total of 18 published sMRI studies were retrieved for the meta-analysis (see Fig. 1 for the flowchart), from which 20 independent samples were included for the main analysis on psychopathy. The main analysis included pooled data from case-control analyses from 12 samples (296 cases versus 325 controls) and whole-brain relationship with severity of psychopathic traits (total score) among 10 samples (675 participants). Two samples included both analytic approaches from which coordinates were pooled for the main analysis (Table 1). Mean age across the 20 samples was 31.8 years (SD=5.34, range 21.1-39.8). From these 18 studies, 12 reported recruiting participants through correctional/probation settings, 1 through forensic settings, and 5 through community. Following our recent work ^46^, preliminary subanalyses were conducted on psychopathy subdimension in which six samples were included for both F1 (6 samples) and F2 (6 samples).

**Fig. 1.**
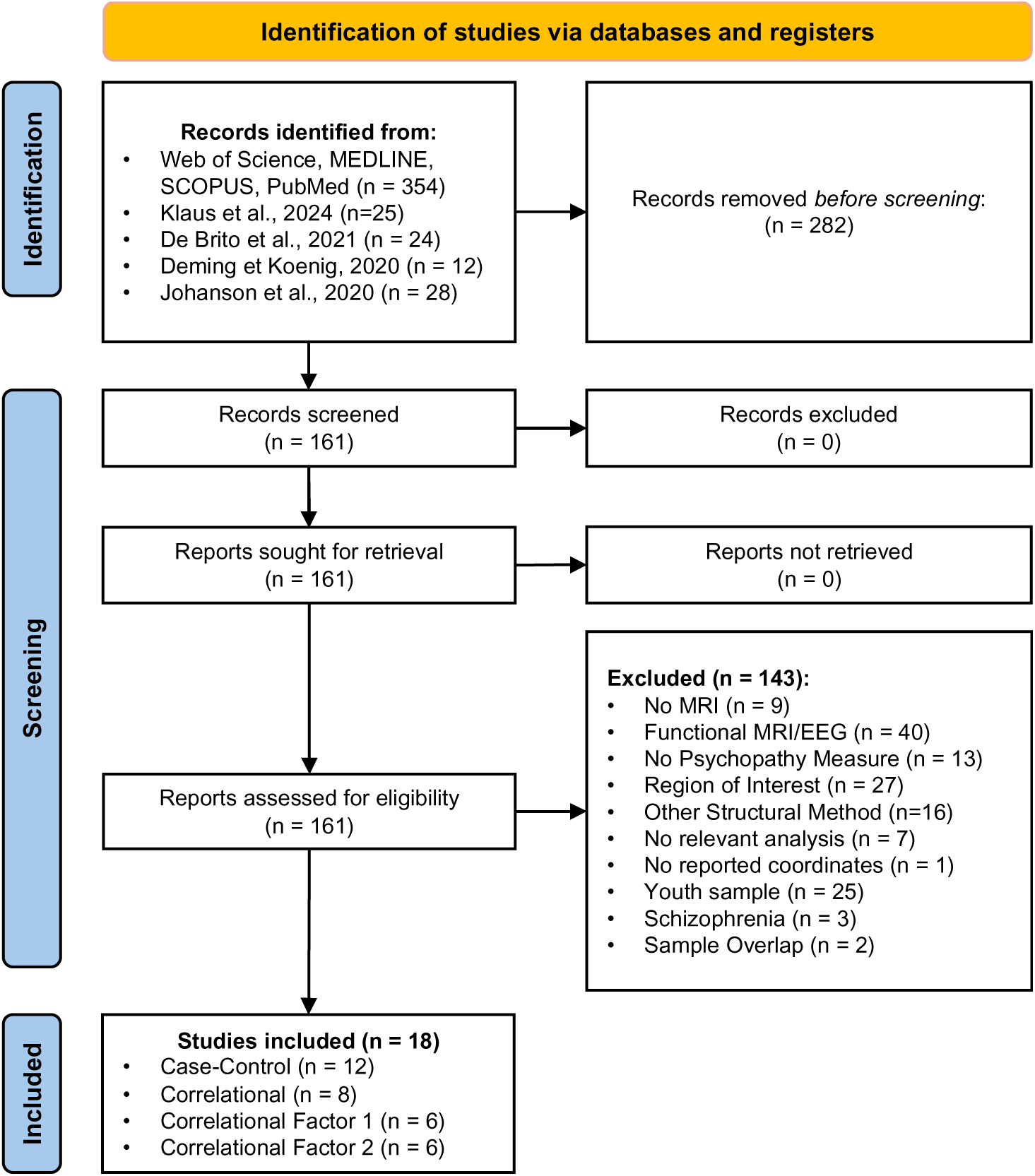
Flowchart of the updated literature search for voxel-based morphometry studies of grey-matter volume on psychopathy after February 2020. Records removed before screening (n=282) included: duplicates (n=195), editorial, reply letter, book chapter & case reports (n=27), literature reviews and meta-analyses (n=46), and unrelated/irrelevant studies (n=14).

**Table 1.** Characteristics of the included Studies (k=20 samples)

| First Author, Year | Sample Characteristics |  |  |  | Psychopathy Assessment |  | Statistical Threshold (Voxel, Cluster Level)<sup>A</sup> |
| --- | --- | --- | --- | --- | --- | --- | --- |
|  | N Cases | N Controls | Mean Age (SD) | Settings | Measures | Total Score |  |
| Beckwith, 2018 (A) <sup>55</sup> | 65 | - | 26.9 (1.1) | Community | PPI | 378.5 | p < 0.005, Cluster Corr |
| Beckwith, 2018 (B) <sup>55</sup> | 90 | - | 26.7 (1.1) | Community | PPI | 355.9 | p < 0.005, Cluster Corr |
| Bertsch, 2013 <sup>56</sup> | 12 | 14 | 27.3 (5.4) | Correctional | PCL-R | 23.8 | p < 0.001, 10 voxels* |
| Chester, 2023 (A) <sup>57</sup> | 53 | - | 27.0 (5.1) | Community | SRP-4 | 50.0 | p < 0.001, 10 voxels* |
| Chester, 2023 (B) <sup>57</sup> | 45 | - | 27.0 (5.7) | Community | SRP-4 | 54.0 | p < 0.001, 10 voxels* |
| Contreras-Rodríguez, 2015 <sup>58</sup> | 22 | 22 | 39.8 (9.2) | Correctional | PCL-R | 27.8 | p < 0.005, 125 voxels |
| Cope, 2012 <sup>59</sup> | 66 | - | 36.9 (7.9) | Correctional | PCL-R | 18.4 | p < 0.005, 5 voxels |
| de Oliveira Souza, 2008 <sup>60</sup> | 15 | 15 | 32.0 (14.0) | Community | PCL-SV | 17.8 | p < 0.005, 5 voxels |
| Ermer, 2012 <sup>61</sup> | 254 | - | 33.9 (9.5) | Correctional | PCL-R | 21.3 | p < .05, Cluster Corr |
| Gregory, 2012 <sup>62</sup> | 17 | 22 | 38.9 (9.4) | Correctional | PCL-R | 28.1 | p < 0.001, 10 voxels* |
| Hofhansel, 2020 <sup>63</sup> | 27 | - | 35.6 (9.5) | Correctional | PCL-R | 16.2 | p < 0.001, Cluster Corr |
| Korponay, 2017 <sup>64</sup> | 41 | 35 | 31.5 (7.7) | Correctional | PCL-R | 32.1 | p < 0.001, Cluster Corr |
| Leutgeb, 2015 <sup>65</sup> | 40 | 37 | 38.1 (12.0) | Correctional | PCL-R | 20.6 | p < 0.001, 10 voxels* |
| Müeller, 2008 <sup>66</sup> | 17 | 17 | 33.0 (5.8) | Forensic | PCL-R | 33.3 | p < 0.001, Not Reported |
| Myznikov, 2024 <sup>67</sup> | 57 | 72 | 24.9 (4.3) | Community | DD-DT | 39.8 | p < 0.001 Cluster Corr |
| Noppari, 2022 <sup>68</sup> | 19 | 19 | 31.0 (7.0) | Correctional | PCL-R | 26.0 | p < 0.001 Cluster Corr |
| Pera-Guardiola, 2016 <sup>69</sup> | 19 | 20 | 39.2 (8.9) | Correctional | PCL-R | 28.1 | p < 0.001, 10 voxels* |
| Schiffer, 2011† <sup>70</sup> | 11 | 27 | 34.55 (6.85) | Correctional | PCL-SV | 13.6 | p < 0.001, 10 voxels* |
| Tiihonen, 2008 <sup>71</sup> | 26 | 25 | 32.1 (8.5) | Correctional | PCL-R | 29.9 | p < 0.005, Cluster Corr |
| Vieira, 2014 <sup>72</sup> | 35 | - | 21.1 (1.8) | Community | TriPM | 71.3 | p < 0.005, ≈144 voxels |

Note. (A, B) refers to independent samples A, B within the same study. <sup>A</sup> = to minimize heterogeneity in statistical thresholding used between studies, threshold was set at p<0.001 with 10 voxel extent (given a 2mm3 voxel), when data were available. \* = raw statistical images were shared and re-thresholded given their respective voxel size. †Data was re-analyzed by conducting a median split on PCL-SV and comparing the above-median group against controls. PCL-SV = Psychopathy Checklist – Short Version <sup>73</sup>; PCL-R = Psychopathy Checklist – Revised <sup>74</sup>; PPI = Psychopathic Personality Inventory <sup>75</sup>; DD-DT = Dirty Dozen / Dark Triad <sup>76</sup>; TriPM = Triarchic Psychopathy Model <sup>77</sup>; SRP-4 = Self-Report Psychopathy, Fourth Edition <sup>78</sup>.

### 2.2. Activation Likelihood Estimation Meta-analysis

A coordinate-based meta-analysis was conducted with the ALE algorithm using peak coordinates of the 20 samples (280 foci). After correction for multiple comparison (cFWE<0.05), a small effect was found in the inferior temporal gyrus (x=-60, y=-20, z=-26, 89 voxels) (see Fig. 2A). More thorough investigation, in which we modeled each peak coordinates with a 4 mm sphere, revealed that the spatial overlap across foci of the 20 samples was very poor (i.e., maximum overlap = 15%; 3 studies) (Fig. 2C). Subanalyses revealed no significant effect when investigating case-control (12 samples, 205 foci) and correlational studies (10 samples, 75 foci), separately. Preliminary analyses on psychopathy subdimensions revealed neither significant effect for F1 (6 samples, 35 foci) and F2 (6 samples, 60 foci).

**Fig. 2.**
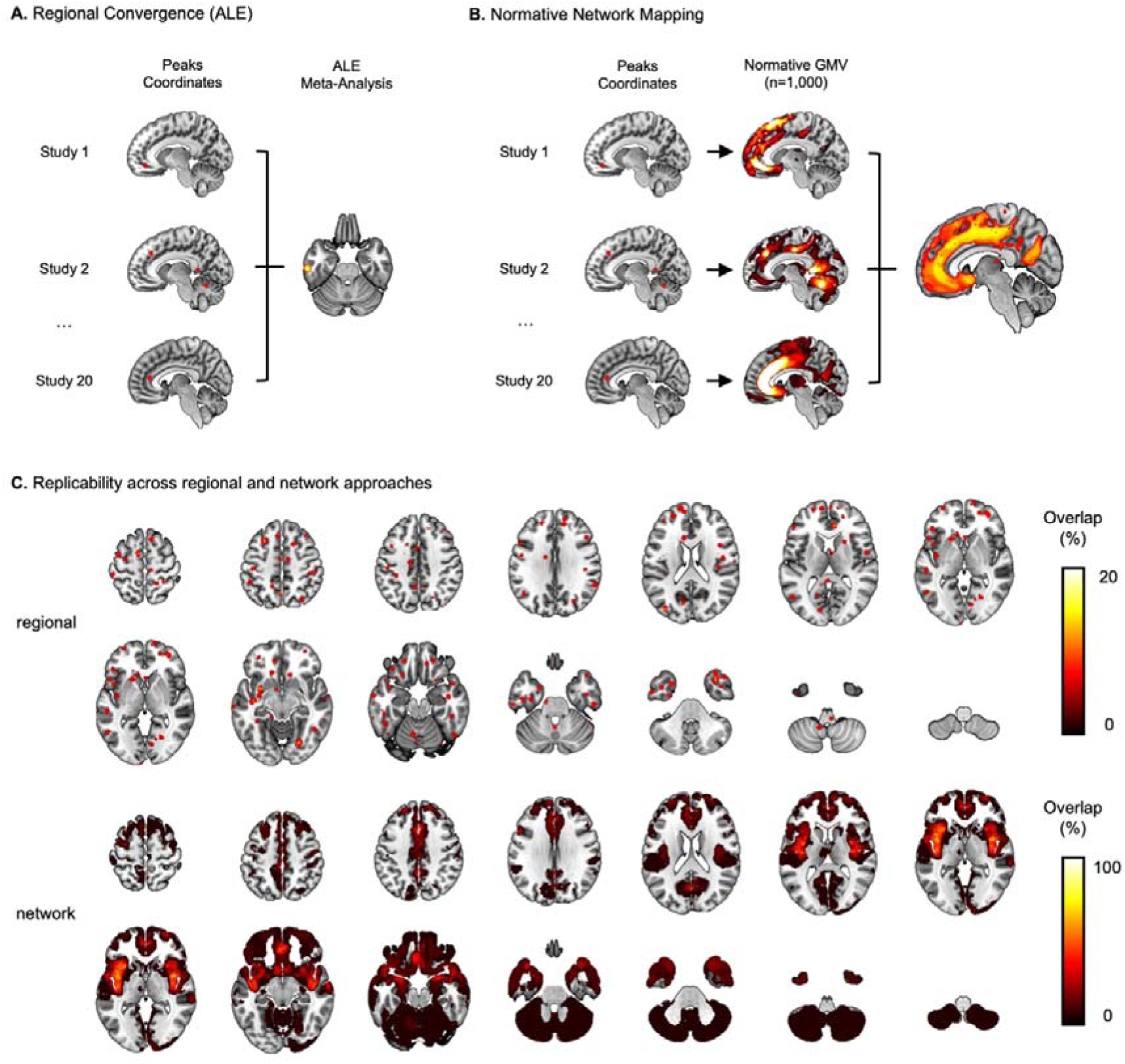
Spatial Convergence across neuroimaging studies on Psychopathy. **A.** Activation Likelihood Estimation meta-analysis was conducted on peak coordinates of 20 VBM samples examining GMV in Psychopathy, which revealed a small cluster in the left inferior temporal gyrus. **B.** Network mapping was subsequently performed to identify a convergent network across the 20 samples. **C.** Displays the differences in overlap between traditional approaches involving spatial convergence based on peak coordinates, and our framework assessing spatial convergence at a network level. The former resulted in low *regional* replicability (up to 15%) (upper row), and the latter produced high *network* replicability between samples (up to 70% of studies)(lower row).

### 2.3. Normative Network Mapping

By modelling structural covariance networks of heterogenous locations, we found significant convergence in many regions including prefrontal, temporal and subcortical structures, after correction for multiple comparisons (TFCE FWE p<0.05). (Fig. 3A). This Structural Network consisted of many key connectivity hubs showing equal or greater replicability than 60%, such as bilateral mid-Insula (70% in both hemispheres), bilateral anterior insula extending to lateral OFC/ventrolateral PFC (60% both hemispheres), posterior midcingulate cortex (60%) (see Supplementary Table 5 for peak coordinates of regions showing ≥ 60% replicability).

**Fig 3.**
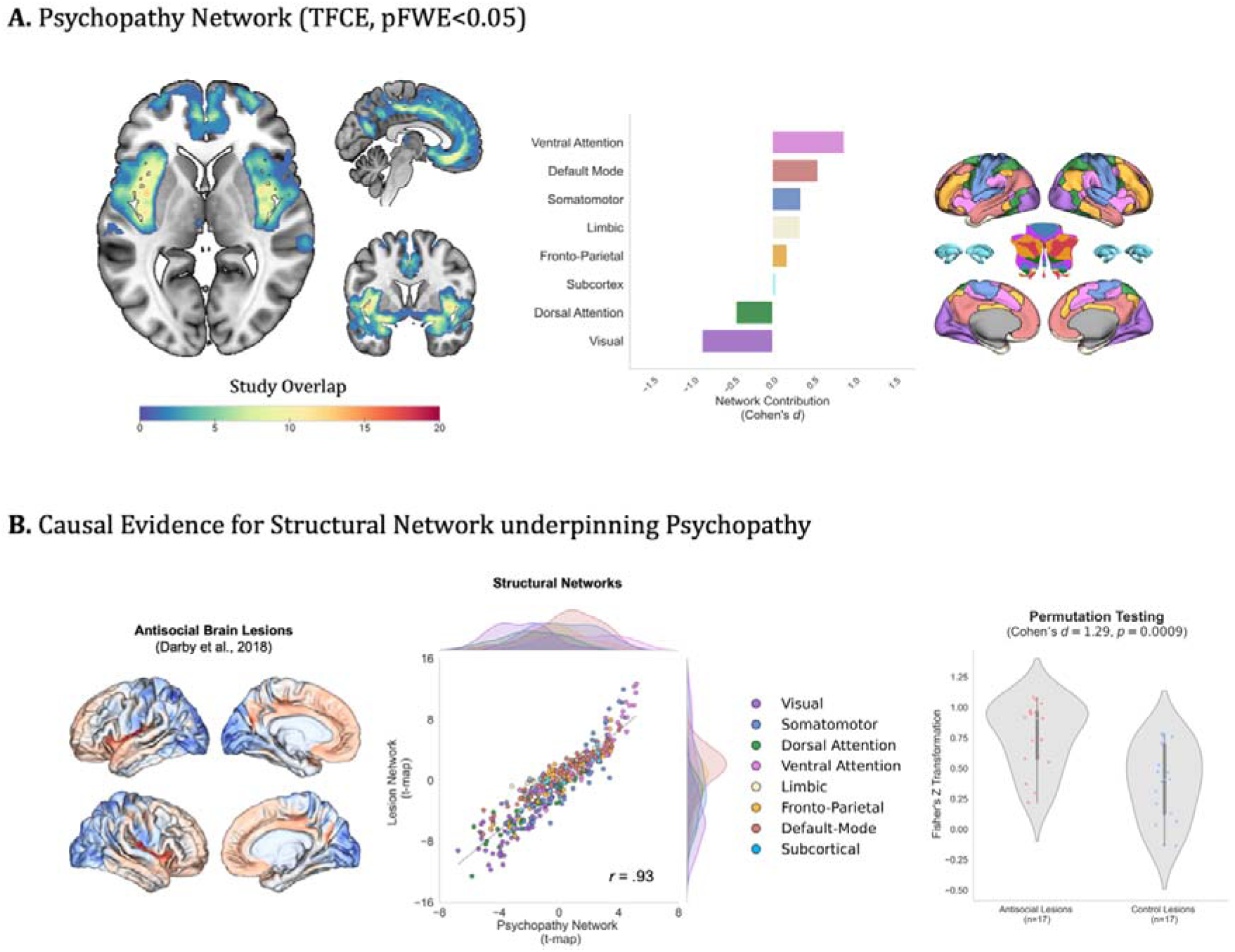
Neural Architecture of the Psychopathy Network. **A.** The Psychopathy Network was formed by the following key hubs: posterior and middle insula extending to superior temporal gyrus, anterior insula extending to lateral OFC/ventrolateral PFC, posterior midcingulate cortex, medial OFC/vmPFC, posterior OFC, and amygdala. This network was primarily driven by stronger structural covariance from nodes of the ventral attention/salience and default mode networks. The bar plot represents the effect size (Cohen’s *d*) of each network’s contribution. **B.** The validity of the Psychopathy Network was assessed by testing its association with a group-level lesion network derived from 17 cases who developed severe antisocial behaviours following brain lesions ^45^ (left, middle panels). A near perfect spatial association was found between our Psychopathy Network and the lesion network (ρ = 0.93, pL<L0.001). This relationship appeared to be stronger when restricting to samples from studies case-control design (ρ = 0.94, pL<L0.001, not shown), compared to samples from studies using a correlational design (ρ = 0.87, pL<L0.001). At the subject level (right panel), the lesion networks of these individuals showed greater spatial similarity to the Psychopathy Network than did 17 matched control lesion networks.

#### 2.3.1. Contribution of Intrinsic Networks

We investigated whether the identified Structural Brain Network may be primarily explained by specific intrinsic connectivity network. Effect sizes (Cohen’s *d*) of Schaefer-400 parcels 7 Networks ^79^ and a subcortical network ^80^ were calculated as a measure of their contribution relative to the rest of the brain. We observed greater effect sizes for nodes of the ventral attention (*d* = 0.88), default mode (*d* = 0.56), followed by somatomotor (*d* = .35) and Limbic (*d* = .33) networks (Fig 3. A right panel). Furthermore, we found similar contribution of intrinsic networks across case-control and correlational samples, and across F1 and F2 (Supplementary Fig. 2).

### 2.4. Multimodal Validation

#### 2.4.1. Brain lesions causally linked to antisocial behaviors

By following the same network mapping approach using 17 masks of cases who developed severe antisocial behaviors after brain lesions ^45^, we found that the lesion-network mapped almost perfectly with the Psychopathy Network when examining it at a group-level (ρ = 0.93) (Fig. 3B middle panel). Moreover, the 17 antisocial brain lesions showed significantly greater similarity with the psychopathy brain network (M = 0.76, SD = .28) compared to 17 matched control lesions (M = 0.38, SD = .31), suggesting specificity of the network to antisocial behaviours (Cohen’s *d* = 1.29, p = 0.0009) (Fig. 3. B right panel)

#### 2.4.2. Cross-Species Validation

To examine how psychopathic traits relate to variations in the identified structural network, we first calculated a participant-specific brain score as a weighted sum of GMV values, using the structural network t-values as weights (Fig 4A).

**Fig 4.**
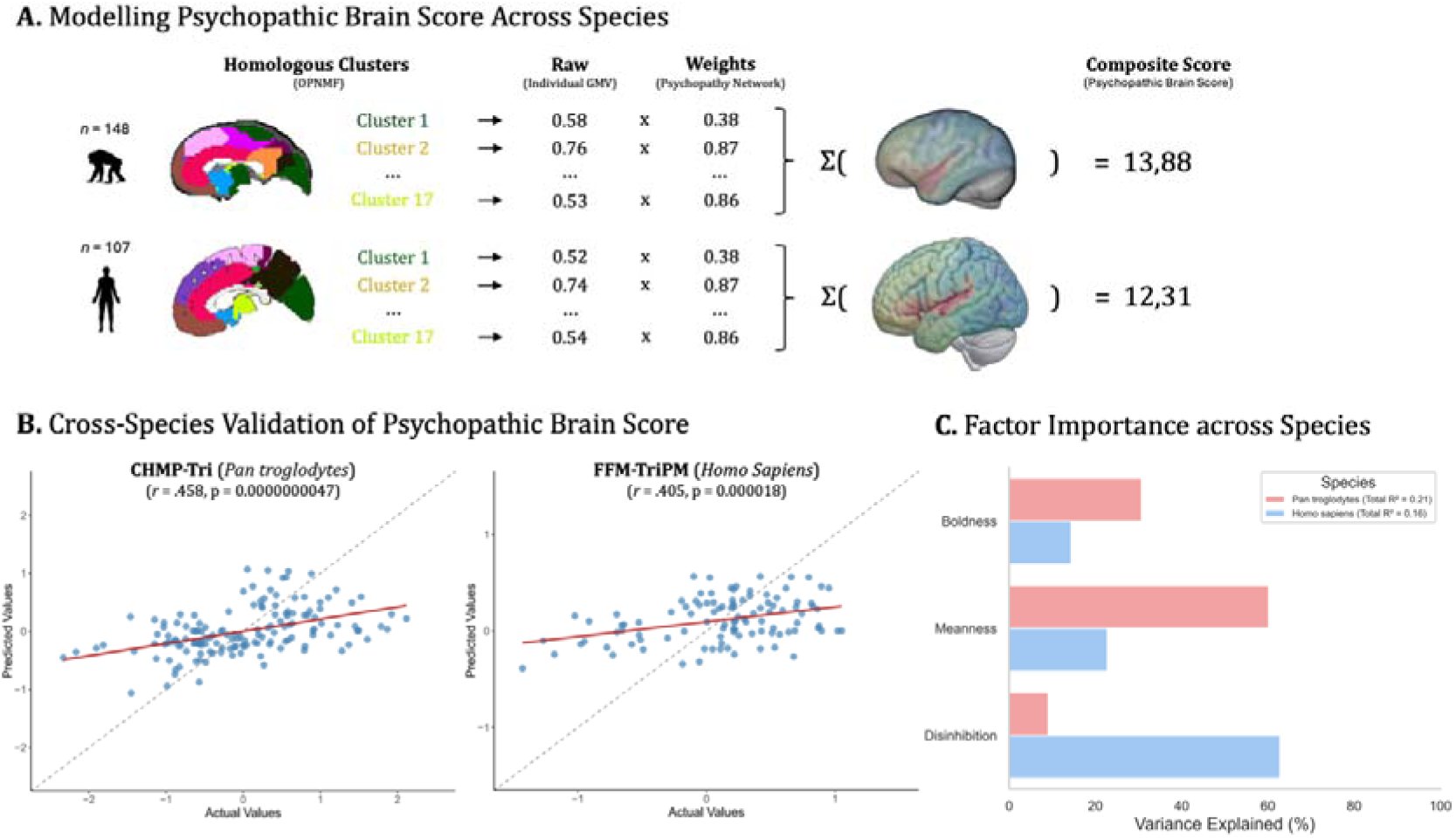
Cross-Species Validation of the Psychopathic Brain Scores. **A.** displays the workflow used to calculate the psychopathic brain scores in chimpanzees (top row) and in humans (bottom row). Briefly, the psychopathy network t-map was parcellated into 17 parcels that were recently identified as being homologous between humans and chimanzees ^81^(left panel). These values were multiplied to raw subject-specific GMVs (middle panel) and summed across the 17 clusters to generate a composite score of psychopathic brain (right panel). **B.** shows the correlation between predicted values and actual values, measuring how well TriPM subscales predicted the unique variance attributed to the Psychopathic brain score, adjusting for TIV. **C.** depicts the results of the dominance analysis, revealing cross-species differences in the contribution of TriPM factors to the prediction of the psychopathic brain score.

In chimpanzees, the TriPM subscales explained a total of 21.01% of the unique variance of the GMV risk scores (*r* = 0.46, p<1.00E-08), after accounting for TIV. Specifically, boldness (standardized dominance *R*^2^ = 30.7%, B = .27, p < 0.001), meanness (standardized dominance *R*^2^ = 60.3%, B = -.41, p<0.001), and disinhibition (standardized dominance *R*^2^ = 9.1%, B = .16, p = .026) were significantly associated with the unique variance of the GMV risk scores (Fig 4. B).

In humans, the Five-Factor composite scores of the TriPM subscales explained a total of 16.4% of the unique variance of the GMV risk scores (*r* = 0.41, *p*<1.00E-04), after accounting for TIV. Specifically, disinhibition (standardized dominance *R*^2^ = 62.8%, B = .41, p = .001) and boldness (standardized dominance *R*^2^ = 14.4%, B = .19, p<0.022), but not meanness (standardized dominance *R*^2^ = 22.8%, B = -.11, p<0.248) were significantly associated with the unique variance of the GMV risk scores (Fig 4. B).

## 3. Discussion

The present study sought to determine, for the first time, whether the apparent poor replicability of GM alterations in psychopathy could be addressed through a network-based framework, beyond traditional meta-analytic approach. We first conducted an updated meta-analysis of 18 VBM studies on psychopathy (20 samples) and observed only a small significant effect in the left inferior temporal gyrus. We further demonstrated that this result was driven by only 3 samples out of 20 independent samples. Crucially, however, we showed that even though peak locations are highly heterogeneous between samples, their structural covariance profile does in fact robustly map onto a common structural network. This network, labelled the Psychopathy Network, was characterized by high replicability of key hubs reaching up to 70% overlap, including the insula (anterior to posterior) which extended to lateral OFC/ventrolateral PFC, and the posterior midcingulate cortex. Validity of this network was first demonstrated by its strong association with a lesion network derived from 17 focal brain lesions causally implicated in the emergence of antisocial behaviours. Finally, we showed for the first time that inter-individual differences in brain regions supporting this network can be predicted by the severity of triarchic psychopathic traits (TriPM) in both humans and chimpanzees, although some discrepancies emerged at the subdimension level. Taken together, our findings highlight the promise of a structural network-based framework in understanding psychopathy, revealing that despite poor replicability of isolated grey matter alterations, these heterogenous findings do in fact converge onto a robust and replicable structural network. The cross-species generalizability, observed in both humans and chimpanzees, suggests that this network may reflect evolutionarily conserved neurobiological substrates of psychopathy.

Decades of research on the structural neuroimaging correlates of psychopathy have highlighted many cortical and subcortical regions known to be implicated in morality and decision-making, especially prefrontal cortices and the amygdala ^82–84^. Despite some recent concerns about the reliability of neuroimaging findings in the field of psychopathy ^27–29^, here we provide clear support for GMV alterations in many brain regions underpinning the paralimbic dysfunction model of psychopathy ^26,85^. More importantly, we show that these paralimbic regions, spanning both the ventral attention/salience and default mode networks, showed a near-perfect association with a structural network of 17 brain lesions that temporally preceded the emergence of antisocial behaviors in a separate sample ^45^. These lesions were structurally connected to a network that was much more similar to the psychopathy network than did 17 matched stroke lesions. In other words, brain lesions known to produce a clinical profile resembling psychopathy, referred to as *pseudopsychopathy* ^18^ or *acquired sociopathy* ^17^, share similar structural underpinnings with psychopathy, as also demonstrates across brain functions ^46^. Given that many of the identified cases with brain lesions committed severe crimes, including murder and sexual aggression ^45^, our findings carry important clinical implications for the prevention of aggressive and violent behaviours. Indeed, psychopathy is well known to be associated with an increased likelihood of institutional violence, intimate partner violence and violent recidivism ^86^. Consequently, patients presenting alterations in GMV and/or brain lesions that structurally covary with the regions identified in our study might need to be identified early and provided with appropriate resources, which in turn could improve the prevention and management of violent behaviours.

Psychopathy is known to be multi-faceted personality disorder, which is commonly characterized by two distinct, yet interrelated dimensions: Interpersonal & Affective, and Chronic Antisocial & Lifestyle dimensions ^4^. Recently, some have expressed their concerns regarding the limited consideration for the heterogeneity psychopathy dimensions in neuroimaging research ^39^ given that these dimensions are linked to distinct psychological and behavioural correlates ^4,87,88^. Interestingly, here we showed that both F1 and F2 appear to rely on a same structural network. GMV differences between F1 and F2 may nevertheless exist. Indeed, across the experiments included in our subanalyses, 69% of the peaks in F1 experiments favoured GMV reduction (i.e., 3 samples reported reductions only, 1 enlargement only, and 2 reported both directions), whereas 65% of the peaks in F2 experiments favoured GMV enlargement (i.e., 4 samples reported enlargements only, 1 reduction only, and 1 both directions). The structural network identified in the present study aligns closely with recently described functional networks ^46^, representing a key step toward understanding the neural architecture underlying psychopathy (Supplementary Fig 3). Although both factors share structural neurobiological features, differences in their functional profiles likely explain their distinct psychological and behavioral correlates ^4,87,88^.

Importantly, we further tested the validity of this structural network across species, examining whether heterogeneous psychopathic traits could predict individuals’ GMV risk scores derived from our network. In both humans and chimpanzees, psychopathic traits (i.e., boldness, meanness, disinhibition) were highly predictive of inter-individual variability in GMV risk scores (16.4% and 21.01%, respectively). Unique variance of meanness was associated with lower brain scores (i.e., smaller volume covariation), whereas the unique variance attributed to boldness and disinhibition predicted greater brain scores (i.e., larger volume covariation). Interestingly, we found cross-species differences in the explanatory power of specific psychopathic traits. Meanness and boldness appeared to play a pivotal role in chimpanzees, while disinhibition was more important in humans. One potential explanation for these differences lies in brain adaptations to evolutionary changes in socio-environmental demands. Notably, the brain regions identified in the current study, commonly implicated in social cognition, are among those that underwent the most significant expansion during human evolution ^81^. Evolutionary pressures toward self-control and norm-following may have driven a reconfiguration in brain connectivity among multimodal association areas ^89^, affecting network strength of functions that rely heavily on the default mode network (i.e., language and social cognition) ^90^. These changes, shifting from salience processing and visual detection to a more internally driven mode of functioning ^90^, likely explained different behavioural expression of the same brain patterns. Other possibilities are measurement issues that could arise from using the same label for distinct phenomena (*jingle fallacy*, ^91^), as shown by a moderate correlation in disinhibition factor score between species ^25^, and/or the use of a five-factor proxy of the TriPM, and/or the limited inter-individual variability in disinhibition scale in chimpanzees given their environment. While more extensive research is needed, our study provides the first evidence that our psychopathy structural network captures inter-individual differences in psychopathic traits across human and chimpanzees.

## 4. Limitations

Several limitations need to be acknowledged. First, VBM studies of GMV vary significantly in terms of demographics, clinical features, MRI protocols, and statistical approaches. Although our network-based approach demonstrates relatively high replicability across samples despite these differences, this heterogeneity may still have led to an underestimation of the true replicability. Second, albeit being preliminary, our subanalyses on psychopathy subdimensions were non-significant and the interpretation of the findings should be approached with caution given the small number of included samples. We encourage neuroimaging researchers to utilize psychopathy facets to gain a more fine-grained understanding of their underlying structural correlates. Third, the traditional coordinate-based meta-analysis showed only weak effects, which may be attributed to limited statistical power, as shown by recent work ^92^. Fourth, we used GMV data of a large sample of healthy individuals (n=1,000, 50% females) to map the heterogeneous brain regions onto a common structural network. It is possible that differences in demographics between the included case samples and the normative sample may have resulted in some variations in network. Consequently, future samples using participant-level data should investigate whether matching demographics yield more precise results. Fifth, we relied on a meta-analytic rather than a mega-analytic approach which could have overestimated the replicability using group-level estimates. International, large-scale collaborative initiatives such as the ENIGMA-Antisocial Behavior Working Group, in which individual-level data are shared, may provide a more precise understanding of the neurobiological substrates of Psychopathy. Sixth, given the absence of TriPM measure for our human sample, we relied on a five-factor composite score of the TriPM. While this statistical approach is a proxy measure of the actual TriPM, future research may seek to replicate our findings using the actual TriPM scale. Seventh, demographics of the human and chimpanzees’ samples used for convergent validity do not match those across GMV samples on psychopathy. Similarly, only the TriPM was used to validate our findings. Future studies should seek to replicate our findings using other measures (e.g., PCL-R, SRP-4, PPI) and/or other populations (e.g., offenders, forensic), before reaching any conclusions about their generalizability.

## 5. Conclusion

Over the past decades, neuroimaging meta-analyses and systematic reviews on the structural correlates of psychopathy have yielded inconsistent findings ^10–13^, raising growing concerns about the reliability of such findings ^27–29^. Here, we show for the first time that heterogeneous peak locations across VBM samples are structurally connected to a common network. We further show for the first time that this network is seemingly identical to the network shared by brain lesions that caused antisocial behaviours. Crucially and strikingly, we demonstrate that our findings are strongly associated with the severity of psychopathic traits in both humans and chimpanzees. In response to the growing scepticism about the reliability of neuroimaging findings, adopting a network-based approach offers promising conceptual and methodological advancements for our understanding of the neurobiology of psychopathy.

## 6. Methods

### 6.1. Activation Likelihood Estimation Meta-analysis

A literature search was performed by reviewing studies included from two recent meta-analyses ^11,13^ and two systematic reviews ^10,93^. In addition, a systematic search strategy was conducted, using four search engines (PubMed, Web of Science, EMBASE, and SCOPUS) from February 2020 to October 31st 2024 to identify relevant studies, extending previous search. The following search terms were used: *psychopathy* OR *psychopathic* AND *neuroimaging* OR *MRI.* JRD screened all titles, abstracts, and full-text studies. Uncertain cases were resolved through discussion with SADB.

Preferred Reporting Items for Systematic Reviews and Meta-Analyses (PRISMA, ^94^; Table S1) guidelines were followed. Studies were included in the current meta-analysis if they: 1) included a sample of adult participants (average >=18 years old); 2) included a case-control analytic approach with groups defined based on a scale assessing psychopathy or correlational analytic approach using a scale measuring total score of psychopathic traits and/or the two main subfactors; 3) included VBM of GMV and 4) reported the peak coordinates of the significant effects across the whole-brain. Raw statistical images were provided by the authors for some of the samples (see ^13^), and were re-thresholded at *p* < .001 uncorrected, following current neuroimaging standards for voxel-level thresholding ^95,96^. A minimal cluster-forming threshold of 10 contiguous voxels (assuming 2 mm³ voxels) was applied to minimize false positives in coordinate-based meta-analyses that do not rely on effect sizes.

Peak coordinates provided in Talairach were then converted onto the MNI standardized space using tal2icbm transformation ^97^. Multiple experiments per study were handled by pooling coordinates to form a sample-level map, to minimize the risk of biases ^98^. The risk of inflating findings through directionality-specific meta-analyses (i.e., conducting meta-analyses for increased and decreased peaks, separately) was handled by pooling both reduction and enlargement to assess spatial convergence across direction. Our meta-analysis of VBM samples, which extend previous work ^11,13^, was conducted using JALE (https://github.com/juaml/JALE), a python package for conducting activation likelihood estimation (ALE) meta-analyses (^30,99^, Supplementary Method, http://www.brainmap.org/ale/). Significance of spatial convergence was established by using a threshold of p<0.001 at voxel-level and FWE-p<0.05 at a cluster-level with 5000 permutations, as recommended ^99,100^.

### 6.2. Normative Network Mapping

A neural network mapping approach was conducted to identify whether heterogeneous peak coordinates reported across VBM samples of GMV alterations in psychopathy may be linked to a common network of structural co-variance in the healthy brain. Briefly, a 4-mm sphere was created around each coordinate from each study to create a sample-level binary mask. Then, using GMV data from 1,000 healthy subjects (ages 18 to 35 years old, 50% females) from the Brain Genomics Superstruct Project (Please refer to Supplementary Methods for MRI data acquisition) ^101,102^, we generated the structural covariance profile of each sample-level mask. This allowed us to identify which voxels covaried with GMV of effects of a given study. Specifically, we extracted the mean volume of each sphere of the sample-level mask in each of the 1,000 healthy individuals and correlated them to the volume of other voxels in the brain. A partial correlation, adjusting for TIV, was conducted to generate a sample-level map, namely the structural covariance network associated to the effects of each study.

To identify, if heterogeneous peaks do indeed map onto a common network, two complimentary analyses were conducted: 1) examining the statistical significance of the network strength across samples; 2) quantifying the degree of overlap across samples. For the first approach, sample-level maps were converted to Fisher’s z scores and combined in a voxel-wise one-sample *t*-test to identify a network that is more consistent than expected by chance ^103^. Statistical significance was set at cluster-based Threshold-Free Cluster Enhancement (TFCE) and Family-Wise Error corrections (FWE TFCE<0.05, 5,000 permutations). The second analysis, aiming to quantify the degree of overlap, involved thresholding sample-level t-maps (T > 5), binarizing, and summing. This threshold has been chosen according to previous samples using a similar approach ^46,104,105^, and because it is fixed and not prone to variations between samples, as with the FDR method.

#### 6.2.1. Contribution of Intrinsic Connectivity Networks

Given the known modular organization of the brain that is also observed through structural covariance ^49,53^, we examined whether the structural brain network underpinning psychopathy may have been driven by nodes from a particular intrinsic network, compared to the rest of the brain. To do so, we used 8 intrinsic connectivity networks, namely the Schaefer-400 7 Networks ^79^ and a subcortical network of 14 regions ^80^. Effect sizes (Cohen’s *d*) were computed by the difference between average t-values a network and average t-value outside the given network divided by the pooled standard deviation.

### 6.3. Multimodal Validation

#### 6.3.1. Antisocial Brain Lesions

Brain lesions provide crucial information for understanding causal etiology of neurological and psychiatric syndromes. In fact, early work showed that patients with focal brain lesions manifested a personality disturbance with a similar clinical profile to psychopathy, sometimes referred to “*acquired sociopathy*” or “*pseudopsychopathy*”. In line with our recent work ^46^, we examined spatial correlation between the discovered Psychopathy structural co-variance network and a brain network of lesions that were causally linked to antisocial behaviours ^45^. To do so, we followed the same approach as described in the main analysis for each of the 17 brain lesions masks of cases that were identified through a systematic literature search ^45^. Both the Psychopathy co-variance Network and the Lesion Network were parcellated using brain regions of the 8 intrinsic networks described above, with 7 additional cerebellar regions from the Buckner Atlas ^106^ (Total of 421 regions). Spearman’s rank order (ρ) correlations were then computed on the extracted mean t-values between both sets of brain regions. We further tested the specificity of the psychopathy network by comparing the 17 lesions associated with antisocial behaviours to 17 matched control lesions from Anatomical Tracings of Lesions After Stroke dataset ^107^. Lesions were matched 1:1 using a nearest neighbor approach from the R package MatchIt ^108^ based on the hemisphere of the lesions (i.e., left, right, or both) and the size of the lesion masks (mean voxel Δ = 28.06 [SD = 45.64]). For each of the lesion networks, a Pearson’s correlation was conducted to assess how similar its co-variance network is to the Psychopathy Structural co-variance Network. Lesion groups were then compared on this similarity index (Fisher’s z transformed correlation coefficient) by a two-sample permutation test (5,000 permutations).

#### 6.3.2. Cross-Species Validation

Neuroanatomical data from both human and chimpanzees were used to test whether the structural brain network found across samples on psychopathy is indeed linked to severity to psychopathic traits. First, human T1-weighted MRI scans from 227 healthy individuals (82 females, age range 20-77 years old) were included publicly available Mind-Brain-Body dataset from the Max Planck Institute for Human and Cognitive and Brain Sciences ^109^ (Please refer to Supplementary Methods for MRI data acquisition). Chimpanzees T1-weighted MRI scans from 149 captive animals (96 females; mean age = 26.11, SD = 10.41 age range 9-54 years old) were provided by the National Chimpanzee Brain Resource (https://www.chimpanzeebrain.org/) and included individuals from the National Center for

Chimpanzee Care at The University of Texas MD Cancer Center (N = 87) and the Emory National Primate Research Center (N = 62) (Please refer to Supplementary Methods for MRI data acquisition). The MRI scanning procedures for chimpanzees at both the NCC and ENPRC were designed to minimize stress for the animals. Data were acquired with ethics approval (#YER-2001206) and were obtained before the 2015 implementation of the US Fish and Wildlife Service and National Institutes of Health regulations governing research with chimpanzees. Both human and chimpanzee data samples were preprocessed using CAT12 ^110^ (Computational Anatomy Toolbox; www.neuro.uni-jena.de/cat/) in SPM12 (Statistical Parametric Mapping; www.fil.ion.ucl.ac.uk/spm/software/spm12/). Chimpanzee data were processed following an established chimpanzee-specific CAT12 pipeline ^111^. Briefly, subject images were linearly registered to the template space, and segmented into the three tissue types, GM, white matter, and cerebrospinal fluid, using a species-specific tissue probability map. The resulting tissue maps were then nonlinearly registered to the population template^112^. The deformation fields were used to modulate GM probability maps that were downsampled (2- and 1.5-mm^3^ resolution) and smoothed at 4- and 6-mm full width half maximum in chimpanzee and human samples, respectively. Quality control was performed for both samples by calculating the sample’s inhomogeneity of GM within CAT12. Modulated GM maps with a mean value greater than 2 standard-deviations were flagged for visual inspection.

Cross-species validation was conducted using the Triarchic Psychopathy Model (TriPM), which includes the dimensions of boldness, meanness, and disinhibition ^113^. For chimpanzees, the Chimpanzee Triarchic (CHMP-Tri) scales were used; these scales represent a validated and homologous measure of the human TriPM ^114^. In humans, composite scores approximating the TriPM scales were computed, given the absence of a direct TriPM measure. This was achieved by summing the products of meta-analytically derived effect sizes relating NEO-Five-Factor Inventory facets ^115^ to TriPM dimensions ^116^ and participant-level raw facet scores. NEO-Five Factor Inventory was available for 108 human participants.

To investigate whether our structural brain network is indeed associated with the severity of psychopathic traits across species, we first derived a global brain measure for each individual. The network was first parcellated into 17 data-driven clusters derived from orthogonal projective non-negative matrix factorization ^117^, which were shown to be homologous between humans and chimpanzees ^81^. Using a similar approach that is typically used to derive polygenic risk scores ^118^ and polyconnectomic scores ^119^, we computed a psychopathy brain score for each participant by applying the network weights of these 17 clusters as multiplicative factors to the GMV values of the corresponding cluster for each individual. These weighted values were then summed to yield a global score reflecting individual variation in GMV across regions linked to the structural network of psychopathy. This approach accounts for individual differences in brain-wide GMV patterns, aligning with the principle of neurobiological equifinality. Multiple linear regression and dominance analyses were subsequently performed to assess the contribution of TriPM subscales to individuals’ psychopathic brain scores, controlling for the residual effect of total intracranial volume. Standardized dominance *R*^2^ values were reported to measure the relative contribution of each TriPM dimension to the total explained variance. A final dataset of 148 chimpanzees and 107 humans was used for further analysis after removing one chimpanzee and one human outlier (Z > 3).

## Acknowledgements

We also would like to thank Dr. Michaël Fox and Dr. Ryan Darby for their generosity in sharing their antisocial lesion masks with us. This study did not receive any specific funding. JRD was supported by a postdoctoral fellowship from the Canadian Institutes of Health Research (MFE-181885). SADB was supported by an Economic and Social Research Council Grant (ES/V003526/1). This work was supported, in part, by NIH grants AG-067419, HD-103490, NS-073134, NS-42867, NS-092988 and NSF-2021711 awarded to WDH. SV was supported by an internally funded post-doctoral grant (Hochschule Bochum: “Nachwuchsförderprogramm 2024”)

## Authors Contributions

J.R.D. performed the systematic search, conducted all analyses and wrote the manuscript. S.A.D.B. assisted with the systematic search, interpreted the results and contributed to manuscript preparation. S.V., R.D.L., W.D.H., and F.H. contributed to chimpanzees’ data collection and aggregation, and contributed to manuscript preparation. All authors revised and approved the final version of the manuscript.

## Data Availability

The chimpanzee sample (www.chimpanzeebrain.org) can be provided by NCBR pending scientific review and a completed material transfer agreement. Request for the chimpanzee data should be submitted to: NCBR. The two human neuroimaging datasets, Brain Genomics Superstruct Project (https://dataverse.harvard.edu/dataverse/GSP) and Mind-Brain-Body dataset from the Max Planck Institute for Human and Cognitive and Brain Sciences (https://fcon_1000.projects.nitrc.org/indi/retro/MPI_LEMON.html), are openly available.

## Code Availability

Coordinate-based meta-analysis was conducted using JALE (https://github.com/juaml/JALE), a python package for conducting ALE (Activation Likelihood Estimation) meta-analyses (^99^, http://www.brainmap.org/ale). Structural Network Mapping was conducted using CAT12 (^110^, Computational Anatomy Toolbox; www.neuro.uni-jena.de/cat/) in SPM12 (Statistical Parametric Mapping; www.fil.ion.ucl.ac.uk/spm/software/spm12/) following similar steps as in functional network mapping (^46,104,120^, https://github.com/nimlab/NMH_Stubbs2023).

