## Supplementary Material for "Mapping the Structural Brain Network of Psychopathy: Convergent Evidence from Humans and Chimpanzees"

Jules Roger Dugré, PhD

Department of Psychology, College of Literature, Science, and the Arts, University of Michigan, Ann Arbor, MI, USA 48109

&

Stéphane Alexandre De Brito, PhD;

Centre for Human Brain Health, University of Birmingham, School of Psychology, Birmingham, UK B15 2TT

### **Supplementary Method****s**

#### Activation Likelihood Estimation

Experiments’ coordinates were used for spatial convergence using the Activation Likelihood Estimate method (GingerALE version 3.0.2, ^1^ http://www.brainmap.org/ale). For each experiment, a 3D gaussian probability distribution was modelled around each coordinate foci, weighted by the number of subjects in the experiment. This method is performed to account for spatial uncertainty due to template and between-subject variance ^2,3^. It also ensures that multiple coordinates from a single experiment does not jointly influence the modeled activation value of a single voxel. The probabilities of all activation foci in an experiment were then combined. Voxel-wise ALE scores arise from the union across all these experiment maps. Consequently, a cluster-level corrected threshold was applied on the voxel-wise image. The size of the supra-threshold clusters was compared against a null distribution of cluster sizes derived from simulation of datasets. We used the following statistical threshold: p<0.001 at voxel-level and FWE-p<0.05 at a cluster-level with 5000 permutations.

#### MRI data acquisition

Brain Genomics Superstruct Project (GSP1000 cohort)

GSP1000 cohort contains a fully 1:1 M:F matched dataset of one thousand human participants ages 18 to 35 years that were scanned with a Siemens Tim Trio 3T scanners (Siemens Healthcare, Erlangen, Germany) at Harvard University and Massachusetts General Hospital using the vendor-supplied 12-channel phased-array head coil. Participants were matched based on age and sex High-resolution (1.2 mm isotropic) multi-echo MPRAGE sequences were acquired for a duration of 2 min 12s, with sagittal acquisition orientation, 144 slices, TR=2200 ms, TE=1.5/3.4/5.2/7.0 ms, FA= 7°, TI=1100 ms, Resolution =1.2×1.2×1.2 mm.

National Chimpanzee Brain Resource

Seventy-six chimpanzees were scanned with a Siemens Trio 3 Tesla scanner (Siemens Medical Solutions USA, Inc, Malvern, Pennsylvania, USA). Most T1w images were collected using a three-dimensional gradient echo sequence with 0.6 × 0.6 × 0.6 resolution (pulse repetition = 2300 ms, echo time = 4.4 ms, number of signals averaged = 3). The remaining 147 chimpanzees were scanned using a 1.5T GE echo-speed Horizon LX MR scanner (GE Medical Systems, Milwaukee, WI), predominantly applying gradient echo sequence with 0.7 × 0.7 × 1.2 resolution (pulse repetition = 19.0 ms, echo time = 8.5 ms, number of signals averaged = 8)

MPI-Leipzig Mind Brain-Body

Two hundred and twenty-seven human participants were scanned with a Siemens Magnetom Verio 3 Tesla scanner (Siemens Healthcare GmbH, Erlangen, Germany) equipped with a 32-channel head coil. MP2RAGE sequences were acquired for a duration of 8 min 22 s, with sagittal acquisition orientation, one 3D volume with 176 slices, TR=5000 ms, TE=2.92 ms, TI1=700 ms, TI2=2500 ms, FA1=4°, FA2=5°, pre-scan normalization, echo spacing=6.9 ms, bandwidth=240 Hz/pixel, FOV=256 mm, voxel size= 1 mm isotropic, GRAPPA acceleration factor 3.

### **Supplementary Tables**

#### **Supplementary Table 1.** PRISMA 2020 Checklist

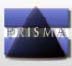
**PRISMA 2020 Checklist**

| **Section and Topic** | **Item #** | **Checklist item** | **Location where item is reported** |
| --- | --- | --- | --- |
| **TITLE** | | |  |
| Title | 1 | Identify the report as a systematic review. | Yes |
| **ABSTRACT** | | |  |
| Abstract | 2 | See the PRISMA 2020 for Abstracts checklist. | S Table2 |
| **INTRODUCTION** | | |  |
| Rationale | 3 | Describe the rationale for the review in the context of existing knowledge. | 2-6 |
| Objectives | 4 | Provide an explicit statement of the objective(s) or question(s) the review addresses. | 6 |
| **METHODS** | | |  |
| Eligibility criteria | 5 | Specify the inclusion and exclusion criteria for the review and how studies were grouped for the syntheses. | 21--22 |
| Information sources | 6 | Specify all databases, registers, websites, organisations, reference lists and other sources searched or consulted to identify studies. Specify the date when each source was last searched or consulted. | 21 |
| Search strategy | 7 | Present the full search strategies for all databases, registers and websites, including any filters and limits used. | 21 |
| Selection process | 8 | Specify the methods used to decide whether a study met the inclusion criteria of the review, including how many reviewers screened each record and each report retrieved, whether they worked independently, and if applicable, details of automation tools used in the process. | 21 |
| Data collection process | 9 | Specify the methods used to collect data from reports, including how many reviewers collected data from each report, whether they worked independently, any processes for obtaining or confirming data from study investigators, and if applicable, details of automation tools used in the process. | NA |
| Data items | 10a | List and define all outcomes for which data were sought. Specify whether all results that were compatible with each outcome domain in each study were sought (e.g. for all measures, time points, analyses), and if not, the methods used to decide which results to collect. | 22 |
|  | 10b | List and define all other variables for which data were sought (e.g. participant and intervention characteristics, funding sources). Describe any assumptions made about any missing or unclear information. | NA |
| Study risk of bias assessment | 11 | Specify the methods used to assess risk of bias in the included studies, including details of the tool(s) used, how many reviewers assessed each study and whether they worked independently, and if applicable, details of automation tools used in the process. | NA |
| Effect measures | 12 | Specify for each outcome the effect measure(s) (e.g. risk ratio, mean difference) used in the synthesis or presentation of results. | NA |
| Synthesis methods | 13a | Describe the processes used to decide which studies were eligible for each synthesis (e.g. tabulating the study intervention characteristics and comparing against the planned groups for each synthesis (item #5)). | NA |
|  | 13b | Describe any methods required to prepare the data for presentation or synthesis, such as handling of missing summary statistics, or data conversions. | NA |
|  | 13c | Describe any methods used to tabulate or visually display results of individual studies and syntheses. | NA |
|  | 13d | Describe any methods used to synthesize results and provide a rationale for the choice(s). If meta-analysis was performed, describe the model(s), method(s) to identify the presence and extent of statistical heterogeneity, and software package(s) used. | 21-22 |
|  | 13e | Describe any methods used to explore possible causes of heterogeneity among study results (e.g. subgroup analysis, meta-regression). | NA |
|  | 13f | Describe any sensitivity analyses conducted to assess robustness of the synthesized results. | NA |
| Reporting bias assessment | 14 | Describe any methods used to assess risk of bias due to missing results in a synthesis (arising from reporting biases). | NA |
| Certainty assessment | 15 | Describe any methods used to assess certainty (or confidence) in the body of evidence for an outcome. | 22 |
| **RESULTS** | | |  |
| Study selection | 16a | Describe the results of the search and selection process, from the number of records identified in the search to the number of studies included in the review, ideally using a flow diagram. | Fig.1 |
|  | 16b | Cite studies that might appear to meet the inclusion criteria, but which were excluded, and explain why they were excluded. | S Table4 |
| Study characteristics | 17 | Cite each included study and present its characteristics. | Table1 |
| Risk of bias in studies | 18 | Present assessments of risk of bias for each included study. | NA |
| Results of individual studies | 19 | For all outcomes, present, for each study: (a) summary statistics for each group (where appropriate) and (b) an effect estimate and its precision (e.g. confidence/credible interval), ideally using structured tables or plots. | NA |
| Results of syntheses | 20a | For each synthesis, briefly summarise the characteristics and risk of bias among contributing studies. | NA |
|  | 20b | Present results of all statistical syntheses conducted. If meta-analysis was done, present for each the summary estimate and its precision (e.g. confidence/credible interval) and measures of statistical heterogeneity. If comparing groups, describe the direction of the effect. | 10 |
|  | 20c | Present results of all investigations of possible causes of heterogeneity among study results. | NA |
|  | 20d | Present results of all sensitivity analyses conducted to assess the robustness of the synthesized results. | 10 |
| Reporting biases | 21 | Present assessments of risk of bias due to missing results (arising from reporting biases) for each synthesis assessed. | NA |
| Certainty of evidence | 22 | Present assessments of certainty (or confidence) in the body of evidence for each outcome assessed. | NA |
| **DISCUSSION** | | |  |
| Discussion | 23a | Provide a general interpretation of the results in the context of other evidence. | 16-19 |
|  | 23b | Discuss any limitations of the evidence included in the review. | 19-20 |
|  | 23c | Discuss any limitations of the review processes used. | 19-20 |
|  | 23d | Discuss implications of the results for practice, policy, and future research. | 21 |
| **OTHER INFORMATION** | | |  |
| Registration and protocol | 24a | Provide registration information for the review, including register name and registration number, or state that the review was not registered. | NA |
|  | 24b | Indicate where the review protocol can be accessed, or state that a protocol was not prepared. | NA |
|  | 24c | Describe and explain any amendments to information provided at registration or in the protocol. | NA |
| Support | 25 | Describe sources of financial or non-financial support for the review, and the role of the funders or sponsors in the review. | 28 |
| Competing interests | 26 | Declare any competing interests of review authors. | 28 |
| Availability of data, code and other materials | 27 | Report which of the following are publicly available and where they can be found: template data collection forms; data extracted from included studies; data used for all analyses; analytic code; any other materials used in the review. | 28 |

*From:* Page MJ, McKenzie JE, Bossuyt PM, Boutron I, Hoffmann TC, Mulrow CD, et al. The PRISMA 2020 statement: an updated guideline for reporting systematic reviews. BMJ 2021;372:n71. doi: 10.1136/bmj.n71. This work is licensed under CC BY 4.0. To view a copy of this license, visit <https://creativecommons.org/licenses/by/4.0/>

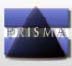
**PRISMA 2020 for Abstracts Checklist**

#### **Supplementary Table 2.** PRISMA 2020 Abstracts Checklist

| **Section and Topic** | **Item #** | **Checklist item** | **Reported (Yes/No)** |
| --- | --- | --- | --- |
| **TITLE** | | |  |
| Title | 1 | Identify the report as a systematic review. | Yes |
| **BACKGROUND** | | |  |
| Objectives | 2 | Provide an explicit statement of the main objective(s) or question(s) the review addresses. | Yes |
| **METHODS** | | |  |
| Eligibility criteria | 3 | Specify the inclusion and exclusion criteria for the review. | No |
| Information sources | 4 | Specify the information sources (e.g. databases, registers) used to identify studies and the date when each was last searched. | No |
| Risk of bias | 5 | Specify the methods used to assess risk of bias in the included studies. | No |
| Synthesis of results | 6 | Specify the methods used to present and synthesise results. | No |
| **RESULTS** | | |  |
| Included studies | 7 | Give the total number of included studies and participants and summarise relevant characteristics of studies. | Yes |
| Synthesis of results | 8 | Present results for main outcomes, preferably indicating the number of included studies and participants for each. If meta-analysis was done, report the summary estimate and confidence/credible interval. If comparing groups, indicate the direction of the effect (i.e. which group is favoured). | Yes |
| **DISCUSSION** | | |  |
| Limitations of evidence | 9 | Provide a brief summary of the limitations of the evidence included in the review (e.g. study risk of bias, inconsistency and imprecision). | No |
| Interpretation | 10 | Provide a general interpretation of the results and important implications. | No |
| **OTHER** | | |  |
| Funding | 11 | Specify the primary source of funding for the review. | NA |
| Registration | 12 | Provide the register name and registration number. | NA |

| **Supplementary Table 3.** Summary of studies included in the current meta-analysis and overlap with prior meta-analyses and systematic reviews | | | | | | | | |
| --- | --- | --- | --- | --- | --- | --- | --- | --- |
| **First Author, Year** | **Included (Case-Control)** | **Included (Voxelwise Correlation)** | | | **Studies overlap** | | | |
|  |  | **Total** | **F1** | **F2** | **Meta-Analyses** | | **Systematic Reviews** | |
|  |  |  |  |  | **De Brito, 2021** | **Deming, 2020** | **Klaus, 2024** | **Johanson, 2021** |
| Beckwith, 2018 (A) ^4^ | - | X | - | - | - | X | X | - |
| Beckwith, 2018 (B)^4^ | - | X | - | - | - | X | X | - |
| Bertsch, 2013 ^5^ | X | - | - | - | X | - | X | X |
| Chester, 2023 (A) ^6^ | - | X | X | X | † | † | - | † |
| Chester, 2023 (B) ^6^ | - | X | X | X | † | † | - | † |
| Contreras-Rodríguez, 2015 ^7^ | X | - | X | X | X | X | - | X |
| Cope, 2012 ^8^ | - | X | X | X | - | X | X | X |
| de Oliveira Souza, 2008 ^9^ | X | - | - | - | X | X | - | X |
| Ermer, 2012 ^10^ | - | X | - | - | - | - | X | X |
| Gregory, 2012 ^11^ | X | - | - | - | X | X | X | X |
| Hofhansel, 2020 ^12^ | - | X | - | X | - | - | X | - |
| Korponay, 2017 ^13^ | X | - | - | - | X | X | - | X |
| Leutgeb, 2015 ^14^ | X | - | X | X | - | - | X | X |
| Müeller, 2008 ^15^ | X | X | X | - | X | X | - | - |
| Myznikov, 2024 ^16^ | X | - | - | - | † | † | † | † |
| Noppari, 2022 ^17^ | X | - | - | - | † | † | - | † |
| Pera-Guardiola, 2016 ^18^ | X | - | - | - | - | X | X | - |
| Schiffer, 2011† ^19^ | X | X | - | - | - | X | - | - |
| Tiihonen, 2008 ^20^ | X | - | - | - | X | - | X | - |
| Vieira, 2014 ^21^ | - | X | - | - | - | - | - | - |
| *Note. † = Study published after the meta-analysis/systematic review.* X = Included; - = Not Included | | | | | | | | |

| **Supplementary Table 4**. Summary of studies excluded from the current meta-analysis but reported by prior meta-analyses and systematic reviews | | | | | | |
| --- | --- | --- | --- | --- | --- | --- |
| **First Author (Year)** | **Reason of Exclusion (Case-Control)** | **Reason of Exclusion (Correlational)** | **Studies overlap** | | | |
|  |  |  | **Meta-Analyses** | | **Systematic Reviews** | |
|  |  |  | **De Brito, 2021** | **Deming, 2020** | **Klaus et al., 2024** | **Johanson, 2021** |
| Kolla, 2014 ^22^ | No comparison with controls | - | - | X | X |  |
| Korponay, 2017b ^23^ | Region of Interest | - | - | - | - | X |
| Ly, 2012 ^24^ | No Voxel-Based Morphometry | - | - | X | - | X |
| Miskovich 2018 ^25^ | - | No Voxel-Based Morphometry | - | X | - | X |
| Nummena, 2021 ^26^ | - | No Coordinates Reported | † | † | X | † |
| Raine, 2000 ^27^ | No Voxel-Based Morphometry | - | - | - | - | X |
| Raine, 2011 ^28^ | No Voxel-Based Morphometry | - | - | - | - | X |
| Sato, 2011 ^29^ | No Statistical Significance | - | - | - | - | X |
| Yang, 2005 ^30^ | No Voxel-Based Morphometry | - | - | - | - | X |
| *Note.* ROI = Region-of-Interest; † = Study published after the meta-analysis/systematic review. X = Included; - = Not Included | | | | | | |

| **Supplementary Table 5**. Summary of Peak Intensities and Replicability within the Psychopathy Structural Network | | | | | | | |
| --- | --- | --- | --- | --- | --- | --- | --- |
| **Regions** | MNI Coordinates | | | **Cluster size (voxels)** |  | Peak Level | |
|  | **x** | **y** | **z** |  |  | **Intensity**  **(*t*-values)** | **Replicability**  **(n, %)** |
| posterior midCingulate Cortex | 2 | -22 | 44 | 31,773 |  | 5.95 | 12 (60.0%) |
| anterior Insula | 46 | 18 | -6 | - |  | 5.75 | 12 (60.0%) |
| anterior Insula | -44 | 16 | -4 | - |  | 5.60 | 12 (60.0%) |
| middle Insula | 40 | 2 | 0 | - |  | 5.29 | 14 (70.0%) |
| dorsal-posterior Insula | 36 | -12 | 8 | - |  | 5.13 | 12 (60.0%) |
| middle Insula | -38 | 0 | 2 | - |  | 5.12 | 14 (70.0%) |
| dorsal-posterior Insula | -38 | -14 | 6 | - |  | 4.46 | 12 (60.0%) |
| Claustrum | -30 | 18 | 0 | - |  | 3.92 | 13 (65.0%) |
| *Note.* Regions were ranked based on their peak intensity. Replicability of these peaks were extracted as a measure of convergence. | | | | | | | |

### **Supplementary Figures**

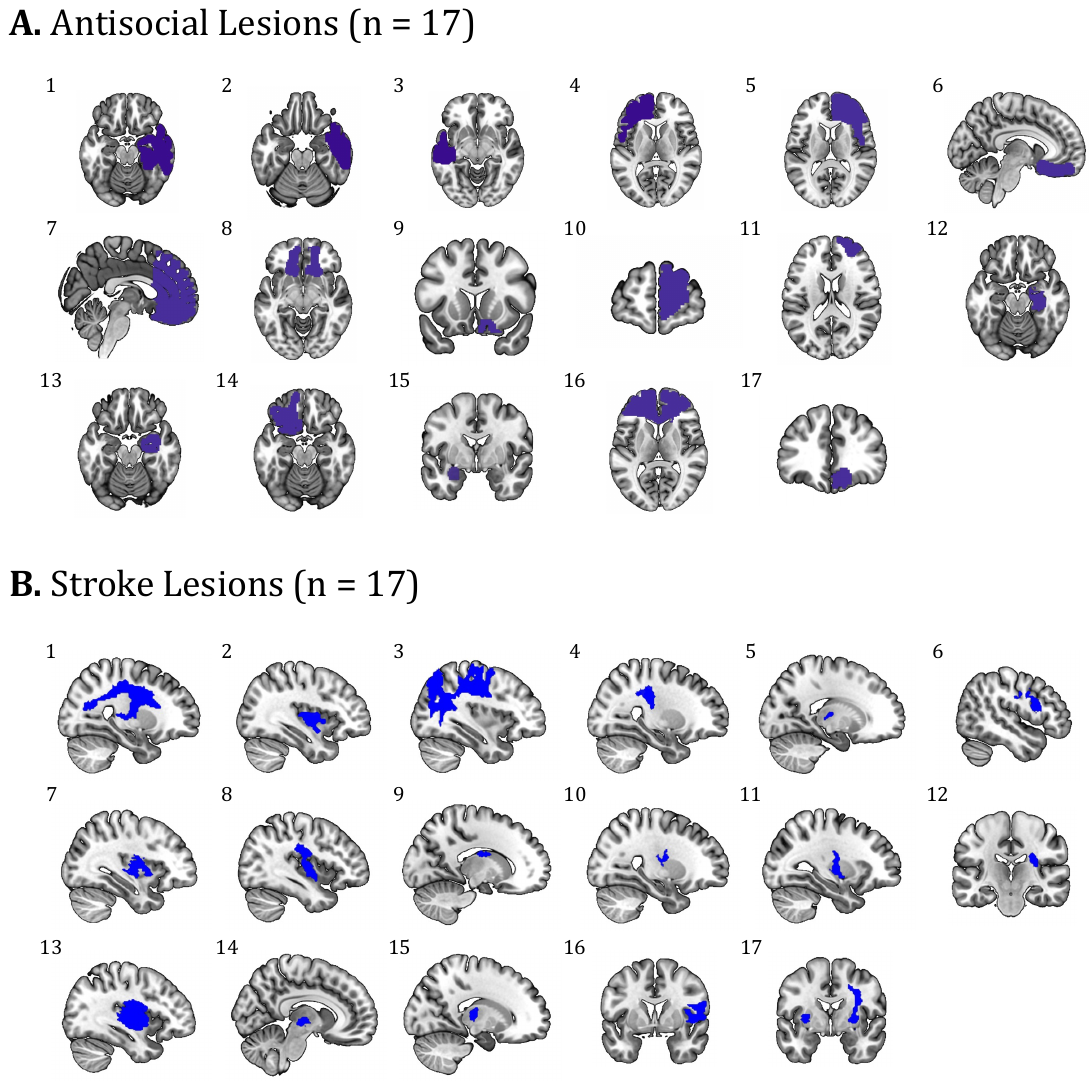

**Supplementary Fig 1.** Lesion masks used in the current study. **A.** displays the 17 antisocial brain lesions (Darby et al., 2018), **B.** shows the 17 stroke lesions that were 1:1 matched in terms of hemisphere location and size.

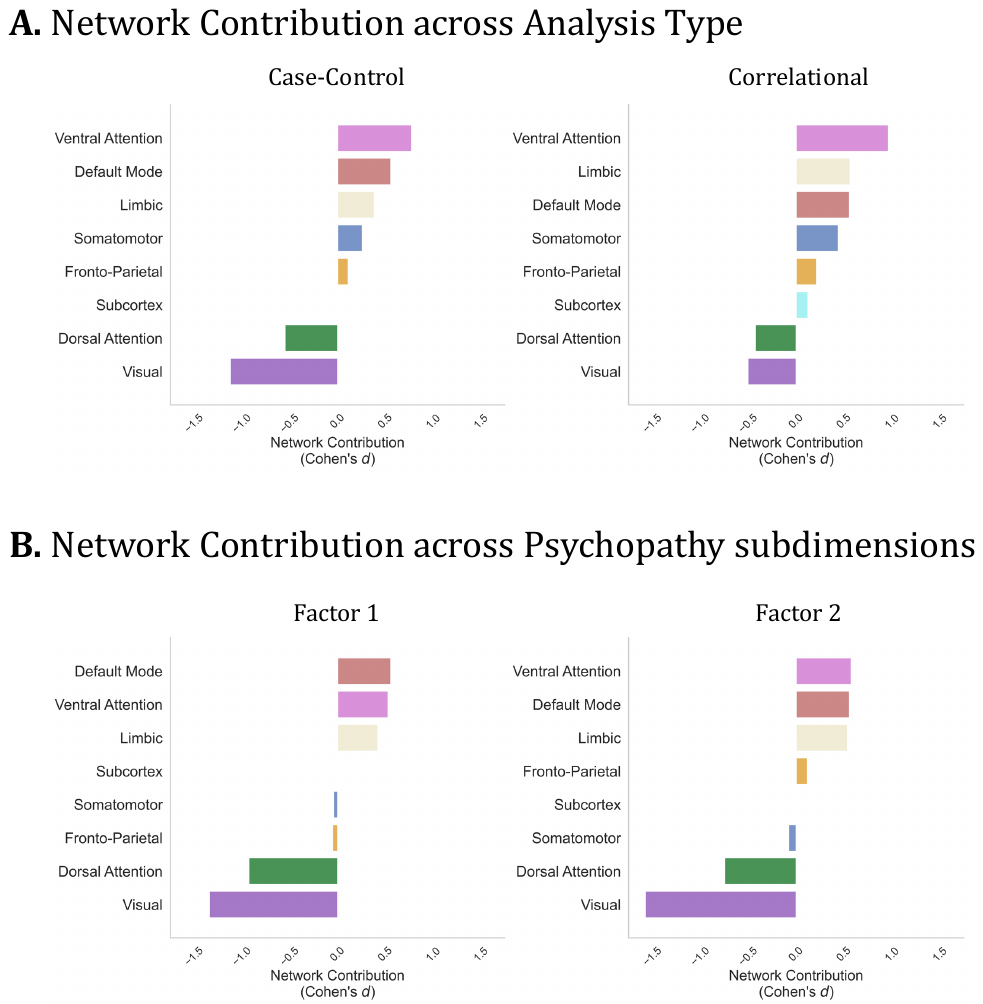

**Supplementary Fig 2.** Subanalyses on Contribution of Instrinsic Connectivity Networks. **A.** displays the effect sizes (Cohen’s *d)* of each of the 8 intrinsic networks in the contributions of a common network among case-control and correlational studies, **B.** displays the effect sizes (Cohen’s *d)* of each of the 8 intrinsic networks in the contributions of a common network among studies on Factor 1, and Factor 2.

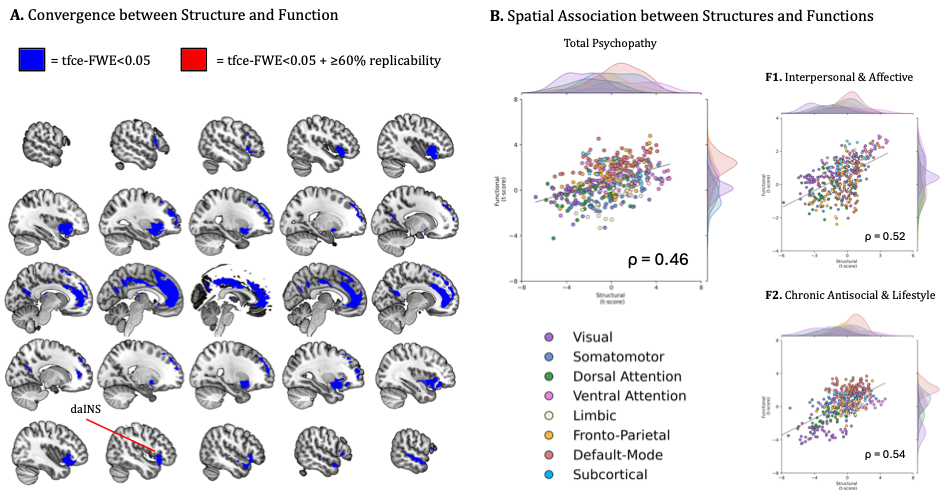

**Supplementary Fig 3.** Spatial overlap between structural and functional networks recently identified by Dugré et De Brito ^31^. **A.** Spatial convergence of brain regions showing high replicability across Psychopathy Structural and Functional Networks (Red = tfce-FEW<0.05; Blue = tfce-FWE<0.05 + >60% replicability). daINS = dorsal anterior Insula; **B.** Spatial correlation between unthresholded t-maps of Psychopathy Structural and Functional Networks (Spearman’s ρ = .46, p<0.001). Association was also found between structure and function for both F1. Interpersonal & Affective (ρ=.52, p<0.001) as well as **F2.** Chronic Antisocial & Lifestyle (ρ=.54, p<0.001). Note. t-maps were parcellated using 421 regions described in the main manuscript.
